# CaMKII Activation Enhances Antioxidant Defence and Mitochondrial Function in human articular chondrocytes

**DOI:** 10.64898/2026.05.27.727824

**Authors:** A Dutta, NJ Day, CS Heluany, D Sochart, BA Fielding, I Smyrnias, G Nalesso

## Abstract

**Background:** The articular cartilage has limited vascular supply and repair capacity, making it particularly susceptible to reactive oxygen species (ROS)-driven oxidative damage. Excess ROS contributes to extracellular matrix breakdown and is a major factor in osteoarthritis (OA) progression. We previously identified Calcium/Calmodulin-dependent protein kinase II (CaMKII) as a regulator of cartilage homeostasis, prompting us to investigate its role during oxidative stress.

**Methods:** Primary adult human articular chondrocytes were isolated from OA cartilage. CaMKII activity was modulated using adenoviral overexpression of a constitutively active form of the kinase or of Autocamtide-2–related inhibitory peptide, a CaMKII inhibitor. Redox status and mitochondrial function were assessed by molecular and metabolic assays.

**Results:** Oxidative stress increased CaMKII phosphorylation. CaMKII inhibition elevated cellular and mitochondrial ROS, whereas CaMKII activation enhanced mitochondrial respiration capacity, improved mitochondrial morphology, and was associated to NRF2 nuclear translocation.

**Conclusion:** CaMKII supports redox homeostasis in chondrocytes and may represent a therapeutic strategy to preserve cartilage integrity and delay disease onset.

## Introduction

OA is a musculoskeletal condition characterised by progressive degeneration of the articular cartilage, synovial inflammation and abnormal remodelling of the subchondral bone, affecting nearly half of the population over the age of 65^1^. Homeostasis in the articular cartilage is maintained by a tightly regulated balance between anabolic and catabolic events regulating the synthesis and degradation of components of the extracellular matrix (ECM).

Currently, only symptomatic treatments are available to manage pain until surgical joint replacement becomes necessary, when pharmacological relief is no longer effective and quality of life is severely compromised.

The development of new, non-invasive, pharmacological treatments for OA is bound to an increase in knowledge of the molecular mechanisms driving its pathogenesis.

We have previously shown that Calcium Calmodulin Kinase II plays a pivotal role in the maintenance of cartilage homeostasis ^2–4^.

CaMKII is a multifunctional serine/threonine kinase playing a crucial role in regulating calcium signalling pathways across various tissues, including the brain, heart, and skeletal muscle. CaMKII is well-known for its involvement in synaptic plasticity, memory formation, and cardiac function ^5–7^.

CaMKII is expressed in 4 different isoforms in mammals, α, β, γ, and δ ^8^. CaMKIIγ and δ are ubiquitously expressed in the human and animal body. We have previously shown that, in human and mouse these two isoforms are expressed in the joint tissues, and γ seems to be quite specifically expressed in the articular cartilage ^4,9^.

Taschner et al and Li et al, showed the importance of CaMKII in mediating chondrocyte differentiation and hypertrophy during development ^10,11^. Previous data showed CaMKII is essential for chondrocyte proliferation and matrix synthesis in response to mechanical stimuli ^12^. Furthermore, Saitta et al. found that CaMKII inhibition in chondrocytes modulates the effects of transforming growth factor-beta (TGFβ) and bone morphogenetic protein (BMP) ^13^.

Our previous work showed that pharmacological inhibition of CaMKII activity can lead to exacerbation of cartilage degeneration in a murine model of OA. Moreover, CaMKII modulation seemed to be associated with upregulation of Heme Oxygenase-1 (HMOX1-1), an enzyme involved the maintenance of the cellular redox balance ^4^. Importantly, oxidative stress has also been observed to modulate the activity of CaMKII in other cell types like cardiomyocytes resulting in atrial fibrillation and fibrosis^14^. These observations suggest that CaMKII may play a role in chondrocyte redox regulation.

Oxidative stress mediated by increased presence/production of reactive oxygen species (ROS) in the joint has been shown to be a key driver of tissue degeneration in OA. ROS are free radicals containing oxygen, such as superoxide and hydroxyl radicals, as well as non-radical molecules like hydrogen peroxide (H₂O₂)^15^. Compounding this oxidative environment, pro-inflammatory cytokines like IL-1β and TNFα induce the expression of genes responsible for nitric oxide and prostaglandin E₂ synthesis, which in turn exacerbate inflammation and contribute to articular tissue damage. Furthermore, excessive ROS can induce DNA damage that drives chondrocyte senescence, which in turn accelerates the progression of the osteoarthritic phenotype^16^.

In this work we aimed therefore to deepen our understanding of the role of CaMKII in the regulation of the chondrocyte’s response to oxidative stress. We induced oxidative stress in primary human chondrocytes and evaluated the effect of CaMKII overactivation and inhibition in the generation of cellular and mitochondrial ROS, mitochondrial activity and activation of an anti-oxidant response. Our results demonstrate that CaMKII inhibition elevates both cellular and mitochondrial ROS. Conversely, CaMKII overexpression promotes NRF2 nuclear translocation, suppresses the activation of the mitochondrial fission protein DRP1, and upregulates the mitophagy marker PARKIN. Our data helps clarifying the importance of this molecule in maintaining the homeostatic balance in chondrocytes and its response to conditions of challenge.

## Materials and Methods

### Cell culture

Adult Human Articular chondrocytes (AHACs) were isolated from preserved regions (Mankin score ≤4)^17^ of cartilage samples obtained from osteoarthritic patients undergoing joint replacement surgery, upon obtainment of informed consent. Samples were provided by the Southwest London Elective Orthopaedic Centre, Epsom, Surrey, UK, upon ethical approval from local authorities (London-Southeast REC, 19/LO/0742). Full-thickness cartilage from femoral heads, condyles, or tibial plateaus was excised aseptically, diced into small cubes, and a portion fixed in paraformaldehyde (PFA) for histology. Articular chondrocytes were isolated and plated as previously described ^2^.

### Adenoviral transduction

To regulate CaMKII activity, recombinant adenoviruses expressing a constitutively active form of CaMKIIγ (AdCaMKII), a peptidic inhibitor of the kinase, the Autocamtide Inhibitory Peptide (AdAIP), and an empty adenoviral vector (AdEmpty) were purchased from VectorBuilder, USA. Chondrocytes were infected at a multiplicity of infection (MOI) of 100 in 2% complete media (CM) (DMEM/F12 + GlutaMAX (1:1) (1X) (Gibco, USA) containing 10% Foetal Bovine Serum (FBS) (Gibco, USA), and 1% antibiotic/antimycotic solution (Thermo Fisher Scientific, USA) for 24 hours. The inoculum was then replaced with fresh 2% CM.

### Western blotting

After treatments, AHAC monolayers (0.4×10^6^ cells per well) were lysed in RIPA buffer (50mM Tris, pH 8 (Sigma Aldrich); 1% Triton X-100 (Sigma Aldrich); 150mM NaCl (Sigma Aldrich); 0.5% sodium deoxycholate (Sigma Aldrich); 0.1% SDS (Sigma Aldrich); containing 10% protease inhibitor (Sigma-Aldrich) and 1% Phosphatase Inhibitor Cocktail 3 (Sigma-Aldrich)). Protein concentrations were determined using the bicinchoninic acid (BCA) assay (Thermo Fisher Scientific). Equal amounts of protein (20–30 µg) were separated by SDS–PAGE using 4–20% Tris–Glycine gels (Thermo Fisher Scientific) and transferred onto nitrocellulose membranes using the Mini Trans-Blot Cell system (Bio-Rad).

Membranes were blocked in protein-free blocking buffer (Thermo Fisher Scientific) and incubated overnight at 4 °C with primary antibodies diluted in the same blocking buffer. CaMKII and phospho-CaMKII were detected using rabbit monoclonal anti-CaMKII (1.75 µg/mL; Abcam) and rabbit monoclonal anti-phospho-CaMKII (0.38 µg/mL; Cell Signalling Technology) antibodies, respectively. DRP1 and phospho-DRP1 were detected using rabbit monoclonal anti-DRP1 (700 µg/mL; Proteintech) and rabbit polyclonal anti-phospho-DRP1 (1 mg/mL; Thermo Fisher Scientific) antibodies. β-actin was detected using a mouse monoclonal anti–β-actin antibody (0.4 µg/mL; Sigma-Aldrich).

### 2′,7′-Dichlorodihydrofluorescein diacetate (DCFDA) assay

AHACs were seeded in 96-well plates (2×10^4^ cells/well), incubated for 24h, washed with PBS, and stained with 20μM 2′,7′-dichlorodihydrofluorescein diacetate acetyl ester (H_2_DCFDA) (Invitrogen)for 45 minutes at 37°C. Cells were treated with H_2_O_2_ (50–600 μM) or adenoviral constructs (AdCaMKII/AdAIP/AdEmpty) for 48h followed by 200μM H_2_O_2_ for 30 minutes. Fluorescence was measured at Ex/Em 485/535 nm using a microplate reader (Tecan). Fluorescence data were background-subtracted and normalised to DAPI intensity to generate fluorescence-per-cell values; fold changes were calculated relative to the control.

### Mitochondrial ROS detection assay

Mitochondrial ROS was detected using MitoROS 580 dye (Abcam). AHACs (3×10^4^ cells/well) were stained with MitoROS dye for 30 minutes at 37°C, washed in PBS, and treated with 200 μM H_2_O_2_ (with/without 5 μM AIP (Sigma Aldrich)). Time-course fluorescence readings (30, 60, 90 minutes) were performed using Ex/Em 540/590 nm (Clariostar). Fluorescence data were background-subtracted and normalised to DAPI intensity to generate fluorescence-per-cell values; fold changes were calculated relative to the control.

### Mitochondrial stress test

Seahorse flux analysis and the mitochondrial stress test (Agilent, UK) were used to assess mitochondrial oxygen consumption rate (OCR) in AHAC. Following appropriate experimental manipulations, AHACs (0.4×10^6^ cells/well) were incubated in assay media (10 mM glucose, 2 mM glutamine, 2 mM pyruvate) for 1 hour, and OCR was measured following sequential injections of oligomycin (2µm), carbonyl cyanide-p-trifluoromethoxyphenylhydrazone (FCCP 2µm), and rotenone/antimycin A (Both 1µm). FCCP concentration was optimised to ensure maximal respiration levels in these cells. Seahorse analysis was performed using the Agilent Seahorse XFe24 Analyzer. OCR values were normalised to protein concentration following quantification of total protein by BCA assay at the end of each experiment. Data are presented as OCR fold changes in AdCaMKII-infected AHACs relative to AdEmpty controls.

### Immunofluorescence

AHACs (30,000/well) were seeded in 8 wells-chamber slides, infected with AdEmpty, AdCaMKII, or AdAIP for 48 hours, and treated with 200μM H_2_O_2_ for 1.5 hours. Cells were fixed in 4% PFA and blocked in Pierce protein-free blocking buffer (Thermo Fisher). AHACs were incubated overnight at 4°C with the primary antibody – anti-Nuclear factor erythroid 2-related factor 2 (NRF2) mouse monoclonal antibody (Santacruz) diluted 1:100 in blocking buffer (Thermofisher, Rockford, USA). The cells were then stained with Alexa Fluor 647-conjugated secondary antibody diluted 1:200 in blocking buffer (Abcam) overnight at 4°C. Upon washing in TBS 0.1% Tween, nuclei were stained with DAPI. Images were acquired via fluorescence confocal microscopy (Nikon A1M Confocal Microscope) with a 40X objective lens, and analysed using CellProfiler.

### Studying mitochondrial morphology

AHACs were infected with AdEmpty or AdCaMKII for 48 hours, treated with 200μM H_2_O_2_ for 1.5 hours, and stained with 200nM Mitotracker (Thermo Fisher). After fixation (4% PFA), autofluorescence quenching with NH_4_Cl, and DAPI staining (18 µM), slides were mounted in Mowiol and imaged using a confocal microscope (Zeiss LSM980) using a 63X objective lens.

### Statistical Analysis

Shapiro-Wilk tests were performed to evaluate the normality of the data. Any outliers in the treatment groups were identified using the ROUT outlier test with a Q value of 1%. Unpaired T tests were performed to evaluate parametric data between two groups. For data sets with multiple groups, analysis of variance (ANOVA) analysis was performed with a Tukey post hoc test to account for multiple comparisons through statistical hypothesis testing. All experiments were performed using independent biological donors, with each experimental condition processed in technical triplicate, and values from the technical replicates were averaged to provide a single representative value per donor for analysis. *P<0.05, **P < 0.01, ***P < 0.0005, ****P < 0.0001, ns = not significant. All the data are presented as mean ± SD.

## Results

### ROS activates CaMKII in AHACs

Oxidative stress has been shown to modulate CaMKII activities in various diseases, including ischemic stroke, cancer, diabetes, cardiovascular disease, and asthma ^6,18–22^. Here, to investigate whether oxidative stress affects CaMKII activation in the articular cartilage, we used H_2_O_2_, as ROS generated from its break-down are known to promote catabolic events linked to OA development and progression ^23,24^.

Firstly, we treated AHACs with different concentrations of H_2_O_2_ (50μM, 200μM, 600μM) for 30 minutes. DCFDA staining was then used to measure intracellular ROS such as superoxide. For each H₂O₂ concentration, cells co-treated with the anti-oxidant N-acetylcysteine (NAC) to confirm the effects observed can be attributed to ROS levels. Phorbol 12-myristate 13-acetate (PMA), a potent inducer of oxidative burst, was included as a positive control. Our results show that increasing concentrations of H_2_O_2_ promoted a dose-dependent increase in ROS generation in the AHACs upon a 30-minute stimulation (Fig 1a).

**Fig 1:**
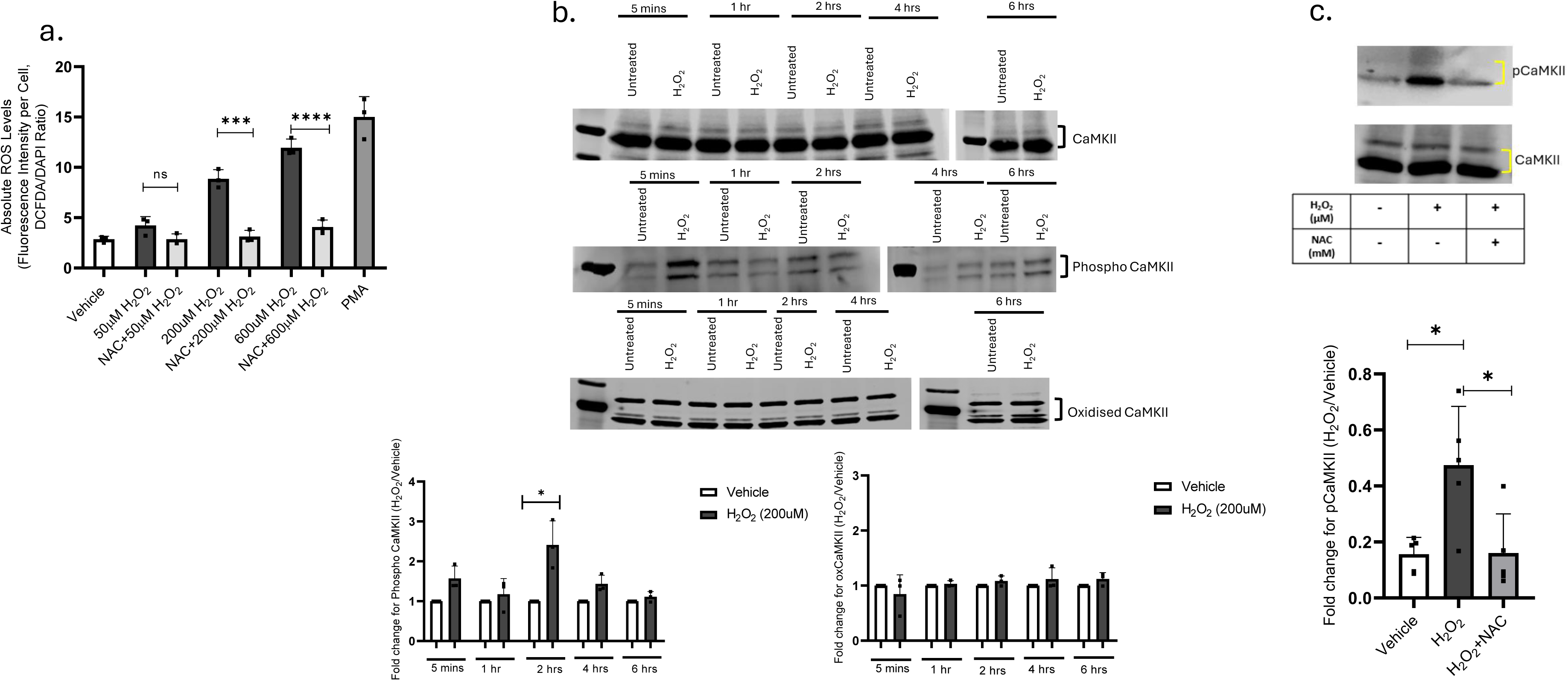
H₂O₂-induced oxidative stress promotes CaMKII phosphorylation. (a) AHACs were pre-stimulated with 10µM NAC and then co-stimulated with 200 µM H₂O₂ alone or in combination with NAC for 5 min, 1, 2, 4 and 6 h. Intracellular ROS were measured through DCFDA staining. PMA at 80ng/mL was used as positive control. Data represent mean ± SD of 3 independent biological replicates. Data analysed with 2-way ANOVA followed by Šidák’s multiple comparison’s post-test. -test (b) Phosphorylation and oxidation of CaMKII (pCaMKII and oxCaMKII) were assessed by Western blotting and quantified as a ratio relative to total CaMKII expression. Data are presented as mean ± SD from 3 biological replicates. Statistical significance between vehicle and H₂O₂ treatment at each timepoint was determined by unpaired t-test(c) AHACs were treated with 200 µM H₂O₂ alone or combined with 10 µM NAC for 2 hours. pCaMKII levels were assessed by western blotting and quantified as ratio relative to total CaMKII expression. Data are presented as mean± SD from N=5 independent biological donors. Statistical significance was determined by One-way ANOVA followed by Tukey’s multiple comparisons test. (N = 5). In all panels vehicle = supplemented EBSS.

Subsequent experiment in AHACS were performed with 200μM H_2_O_2_ over a time course. The expression levels of total oxidized and phosphorylated CaMKII were then measured by western blotting. The immunoblot analysis showed a marked increase in CaMKII phosphorylation at 2 hours following H_2_O_2_ stimulation, however no change in oxidised CaMKII was detected (Fig 1b).

To further validate whether CaMKII phosphorylation was specifically a result of oxidative stress or a consequence of other biochemical processes occurring in AHACs in response to H_2_O_2_ stimulation, we repeated the experiment in the presence of an antioxidant, N-acetylcysteine (NAC).

Co-treatment with NAC and H_2_O_2_ prevented CaMKII phosphorylation induced by the latter (Fig 1c) demonstrating that the H_2_O_2_-mediated phosphorylation of CaMKII is dependent on production of ROS.

### CaMKII modulates chondrocyte response to oxidative Stress

To better understand whether the ROS-induced CaMKII phosphorylation was linked to a pro-anabolic or pro-catabolic activity of the kinase, we inhibited CaMKII by infecting AHACs with an adenovirus expressing the CaMKII peptide inhibitor AIP ^25^ or an empty adenoviral vector as negative control and measured ROS generation by DCFDA staining assay.

Inhibition of CaMKII with AIP exacerbated ROS production in H_2_O_2_ stimulated cells in comparison to control cells. This suggests that CaMKII inhibition leads to enhanced ROS generation, thereby promoting oxidative stress (Fig 2a).

**Fig 2:**
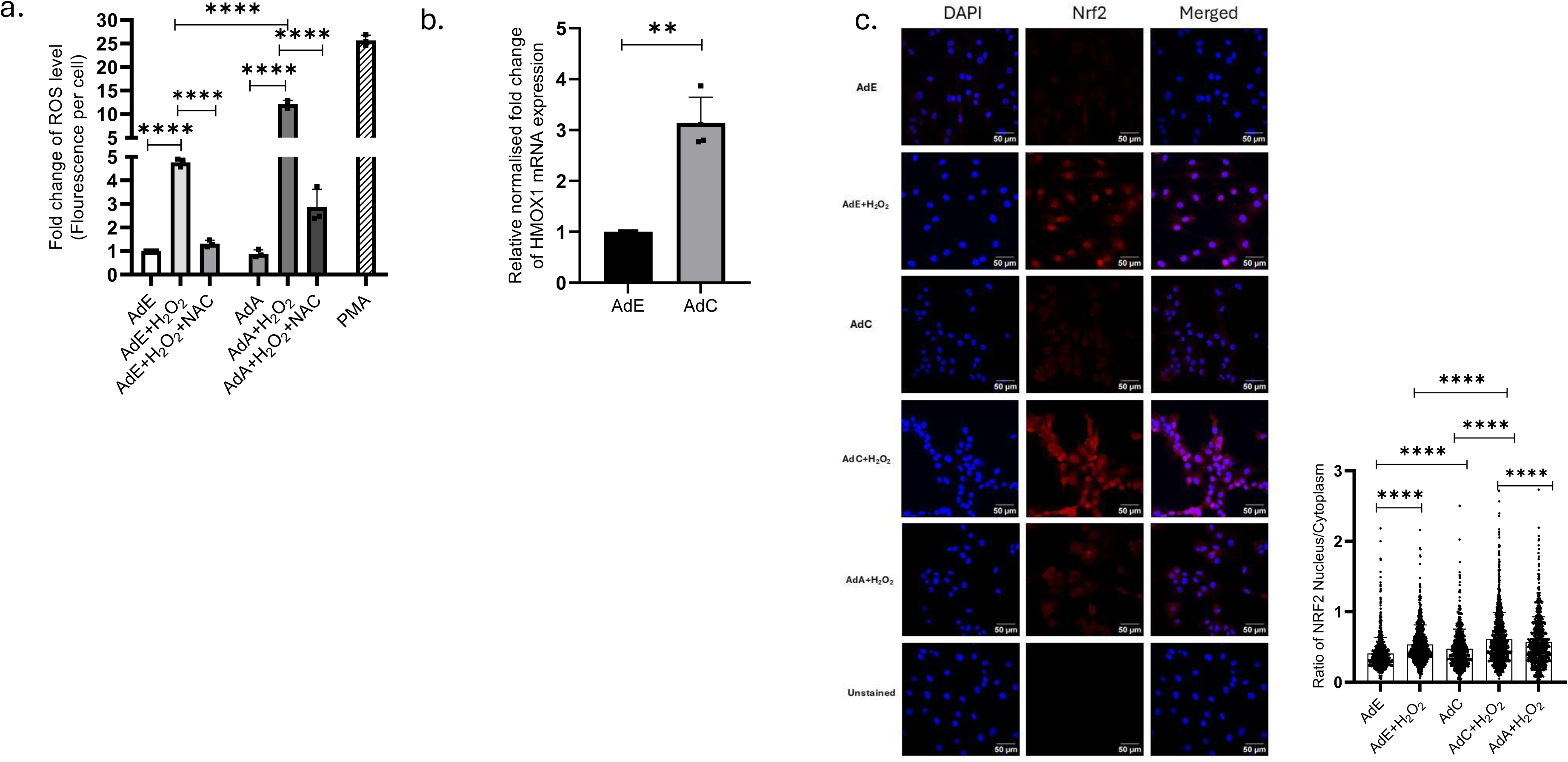
CaMKII modulates chondrocyte response to OS. a. AHACs were transduced with AdA and AdE respectively for 48 hours, followed by H_2_O_2_ treatment for 30 mins (n=3 biological donors). Fluorescence per cell was normalized using DAPI nuclear staining, and fold changes were calculated relative to the control. Statistical analysis was conducted using two-way ANOVA and Tukey post-test. Individual data points represent biological replicates. b. AHACs were infected with AdE and AdC adenovirus for 48 hours. HMOX1 expression was measured by qPCR. All data points have been normalised to 18s expression. Data is presented as the fold change of CaMKII overexpressed (AdC) to control (AdE). The data was analysed using t test. Individual data points represent biological replicates (n=4 biological replicates). **=P<0.005. c. Representative images of Nrf2 translocation to the nucleus in AHACs. Cells were infected with AdE, AdC or AdA virus for 48 hours and then treated with H_2_O_2_ for 1.5 hours. AHACs in the negative control were treated in the same manner except the primary antibody was not added. The intensity of Nrf2 staining was quantified in CellProfiler. Data presented as violin plots where each individual data point represents fluorescent intensity of one cell. The data is presented as the ratio of Nrf2 fluorescence in the nucleus /cytoplasm (N=3 biological donors). 5 random fields per technical replicate per condition for each biological replicate were analysed. Scale bar-50μm. The data was analysed using two-way Anova and Tukey post-test.

Conversely, infection of AHACs with an adenovirus expressing a constitutively active form of CaMKIIγ upregulated the mRNA levels of Heme oxygenase 1 (HMOX1), an NRF2-regulated modulator of the cellular anti-oxidant response ^26^ (Fig 2b).

The NRF2 transcription factor, which mediates antioxidant defence by translocating to the nucleus during conditions of stress to activate a plethora of detoxifying and anti-oxidant genes^27^. To investigate whether the CaMKII-dependent increase in HMOX1 expression (Fig. 2b) might be mediated through NRF2 activation, we examined NRF2 localisation in AHACs. Inhibition of CaMKII with AdAIP markedly reduced the H₂O₂-induced nuclear translocation of NRF2 (Fig. 2c), indicating that CaMKII activity is required for efficient NRF2 activation and suggesting that the upregulation of HMOX1 observed in Fig. 2b might be driven through this pathway.

### CaMKII modulates mitochondrial ROS generation and improves mitochondrial respiration in AHACs

Mitochondria play a crucial role in modulating oxidative stress by regulating ROS production and activating antioxidant defence mechanisms within cells ^28^. CaMKII has been shown to modulate mitochondrial activity in multiple biological contexts, including muscle cells and hepatocytes ^29–31^.

To test whether CaMKII influences mitochondrial ROS production in articular chondrocytes, we measured mitochondrial superoxide, a primary ROS, in the absence of CaMKII using the fluorescent probe MitoSOX Red (Fig. 3a). Mitochondrial ROS levels increased in AHACs in response to H₂O₂ stimulation, which was exacerbated following inhibition of CaMKII with soluble AIP (Fig 3a). These findings indicate a protective role for CaMKII in regulating superoxide generation and maintaining mitochondrial function(s) under conditions of oxidative stress.

**Fig 3:**
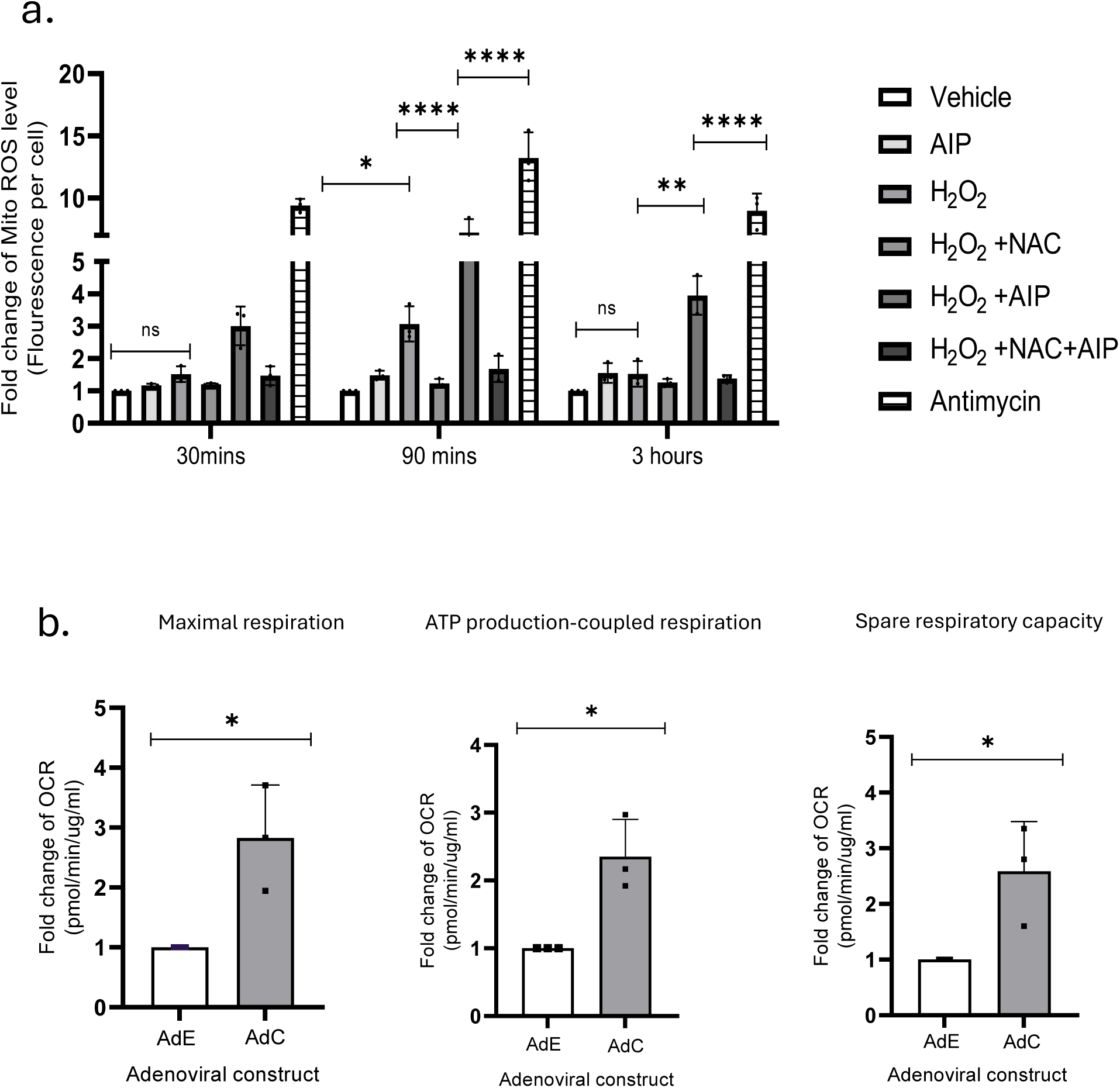
CaMKII modulates mitochondrial ROS generation and upregulates the respiratory potential of the mitochondria. a. AHACs were co-treated with 5 µM AIP and 200 µM H_2_O_2_ or 200 µM H_2_O_2_ alone for 30 mins, 1.5 hours, and 3 hours. The cells were then stained with mitoSOX stain for 30 mins. Fluorescence intensity was measured at 590nm. Fluorescence per cell was normalized using DAPI nuclear staining, and fold changes were calculated relative to control (n=3 biological replicates). Statistical analysis was conducted using two-way ANOVA and Tukey post-test. Each individual data point is a biological replicate. B. ATP coupled respiration of the AHACs, maximum respiration of the AHACs, spare respiratory capacity measured by SeaHorse analysis in AHACs infected with AdE and AdC for 48 hours. Data were normalised to protein concentration measured by BCA assay. The data is represented as the fold change of the OCR of the AdC infected AHACs to the control (AdE infected AHACs). The data was analysed using unpaired t test and (N=3 biological donors).

Dysfunction of the mitochondrial electron transport chain has been reported in OA-affected chondrocytes ^32^. To assess the role of CaMKII in mitochondrial respiration, we measured mitochondrial OCR in AHACs with and without overexpression of CamKII. As shown in figure 3b, overexpression of CamKII with AdCamKII resulted in significantly increased ATP production-coupled respiration, maximal respiration, and spare respiratory capacity as compared to control cells (AdE). Our data suggest that overexpression of CaMKII significantly improves mitochondrial functions, suggesting possible increased resilience of AHACs to oxidative stress. Together, these findings underscore CaMKII’s role in supporting mitochondrial function by modulating superoxide production and optimizing respiratory efficiency.

### CaMKII modulates mitochondrial morphology by regulating mitochondrial fission under oxidative stress

Elevated ROS levels lead to mitochondrial fragmentation, which serves as a cellular response to ROS, allowing damaged mitochondria to be segregated from healthy ones to preserve overall mitochondrial health ^33^. Treatment of AHACs with 200 µM H₂O₂ caused mitochondrial fragmentation which was rescued by overexpression of activated CaMKII (Fig 4a). Then, to understand whether this this effect was mediated through the regulation of mitochondrial fission, we measured the activation of the mitochondrial fission marker DRP1. Phospho-DRP1 levels, significantly increased in cells treated with H_2_O_2_ (200 μM) compared to control Ad AHACs, but overexpression of activated CaMKII with H₂O₂ markedly reduced it (Fig 4b), suggesting that CaMKII may play a role in facilitating the recovery of mitochondrial structure by either reducing mitochondrial fission or promoting the clearance of damaged mitochondria.

**Fig 4:**
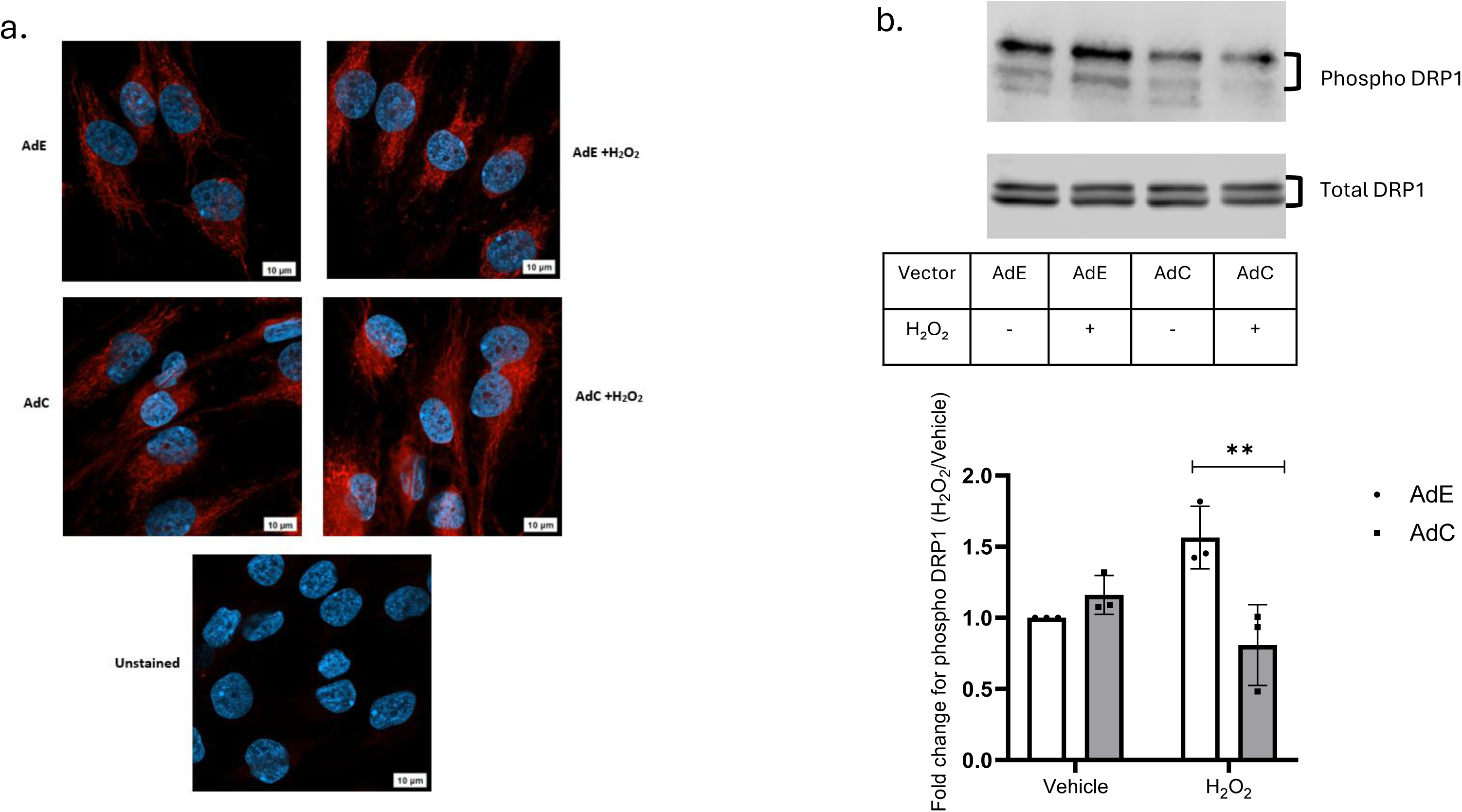
CaMKII modulates mitochondrial morphology by regulating mitochondrial fission under oxidative stress. a. Representative images of mitochondrial morphology affected by CaMKII. AHACs were infected with AdE and AdC for 48 hours and treated with 200uM H_2_O_2_ for 1.5 hours. The cells were then stained with Mitotracker stain and DAPI nuclear stain. AHACs in the negative control were processed in the same way except the cells were not stained with Mitotracker (n=3 biological donors). Scale bar-10μm. b. AHACs were infected with AdE and AdC, for 48h and then stimulated with H₂O₂ or supplemented EBSS as vehicle for 1.5 hours. The phosphorylated form of DRP1 (Phospho-DRP1) was quantified relative to total DRP1 expression, and fold changes were calculated relative to the control (vehicle). The data was analysed with two-way Anova and Tukey post-test. Individual data points represent biological replicates. (n=3 biological replicates). *=P<0.05, **=P<0.01.

### CaMKII activation promotes phosphorylation of PARKIN, an important activator of mitophagy

Mitophagy is a physiological process activated by cells to remove and recycle damaged mitochondria^34^. Our results show that activation of CaMKII in conditions of challenge can contribute to the recovery of mitochondrial morphology, accompanied by a reduction in mitochondrial fission. To understand whether CaMKII could support mitochondrial homeostasis by also aiding in the clearance of damaged organelles, we tested whether overexpression of CaMKII could modulate phosphorylation of Parkin 1 (PARK1) an E3 ubiquitin ligase that accumulates on the outer mitochondrial membrane, thus ubiquitinating outer mitochondrial membrane proteins promoting mitophagy ^35^.

To this end, AHACs were stimulated with 200μM H_2_O_2_ after been infected with AdCaMKII to overexpress CaMKII, or Ad Empty control. Control cells displayed the highest expression levels of total PARK1, representing the basal pool available under homeostatic conditions. In contrast, cells treated with H₂O₂, alone or in combination with AdCaMKII, showed significant PARK1 downregulation. However, downregulation of total PARK1 was accompanied by significantly upregulated levels of phosphor-PARK1 (pPARK1) only in cells infected with AdCaMKII (Fig 5). This data suggest that CaMKII activation contributes to mitophagy induction in response to oxidative stress in articular cartilage.

**Fig 5:**
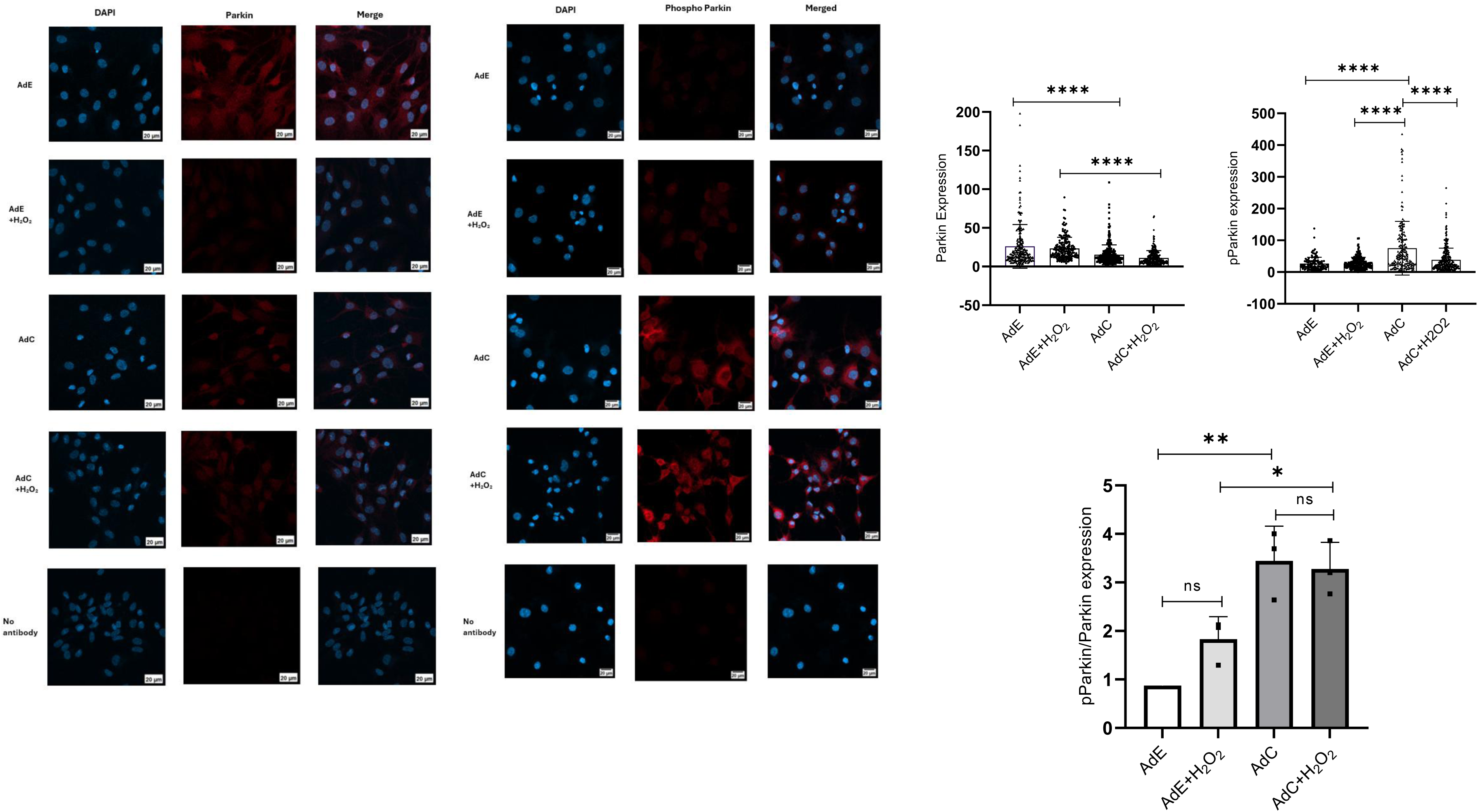
CaMKII activation promotes phosphorylation of PARKIN, an important activator of mitophagy. AHACs were infected with AdE and AdC for 48 hours and then treated with 200uM H₂O₂ for 1.5 hours. Cells were then stained with anti Parkin antibody and anti phospho Parkin antibody. The intensity of Parkin and phospho Parkin staining was quantified by using CellProfiler. The data was analysed using two-way Anova and Tukey post-test. Individual data points represent fluorescent intensity per cell (n=3 biological donors). Scale bar-20μm.

## Discussion

In this study, we demonstrated that CaMKII acts as a key regulator of oxidative stress responses in AHACs, modulating intracellular ROS levels, mitochondrial function, and mitophagy. These findings expand upon previous work suggesting that CaMKII is essential in maintaining cartilage homeostasis^4,13^.

We observed that oxidative stress induced by H₂O₂ increased phosphorylation; but not oxidation of CaMKII. While CaMKII oxidation has been documented in cardiomyocytes ^18^, its absence in our system may reflect tissue-specific differences in redox regulation or methodological constraints. Oxidized proteins, including CaMKII, are often transient and unstable, complicating detection by standard immunoblotting ^36^. Thus, while our data support phosphorylation as the dominant regulatory mechanism in chondrocytes, advanced redox proteomic approaches (e.g., thiol-reactive probes, mass spectrometry) will be needed to more sensitively assess oxidation states in future work ^37,38^.

Our results further indicate that CaMKII activation protects chondrocytes from oxidative stress by enhancing antioxidant responses via NRF2 signalling, thus potentially limiting mitochondrial ROS. This aligns with reports showing that NRF2 activity is impaired in aged and OA cartilage, leading to reduced expression of NRF2-regulated protective enzymes such as HMOX-1 and NADPH quinone dehydrogenase 1 (NQO1) ^39–41^. In animal models, NRF2 activation suppresses inflammation and cartilage damage, underscoring its therapeutic potential ^42^. Interestingly, our findings contrast with prior observations that CaMKII inhibition upregulates HMOX-1 ^4^. Here we show that AIP treatment increases ROS levels, suggesting that HMOX-1 induction under CaMKII inhibition likely reflects a compensatory stress response rather than direct transcriptional regulation. While we focused on HMOX-1, further exploration of additional NRF2 target genes (e.g., NQO1, GCLC, SOD2) will be needed to establish a more comprehensive picture of CaMKII–NRF2 crosstalk.

With respect to mitochondrial function, our data show that CaMKII activation improves ATP production-coupled respiration, maximal respiration, and spare respiratory capacity compared to control conditions, while its inhibition exacerbates ROS production. These findings support the view that mitochondrial respiration is active and physiologically important in articular chondrocytes, consistent with early work demonstrating impaired complex II/III activity, reduced membrane potential, and increased mitochondrial damage in OA chondrocytes ^43^. Mechanistic studies have also shown that while healthy chondrocytes can enhance oxidative phosphorylation under stress, OA chondrocytes display reduced respiratory capacity and shift toward glycolysis ^44^. Nonetheless, CaMKII’s role in regulating mitochondrial metabolism appears to be highly context-dependent. In cardiomyocytes, CaMKIIδ drives maladaptive metabolic remodelling in heart failure and post-infarction cardiomyopathy ^45,46^, while knockout studies suggest CaMKII is dispensable for basal respiration under non-stress conditions ^47^. In the liver, CaMKII even sustains oxidative metabolism during impaired MCU function ^48^. These divergent outcomes likely reflect cell-specific metabolic demands. It is important to note that in our Seahorse assay, the oligomycin-induced decrease in OCR reflects mitochondrial respiration coupled to ATP synthesis. This provides an estimate of ATP-linked respiration but does not quantify total cellular ATP levels. Additionally, the use of high-glucose medium may influence substrate preference. Future studies incorporating varied nutrient environments and direct ATP quantification will be required to more precisely define how CaMKII integrates into chondrocyte bioenergetics.

We also found that CaMKII preserves mitochondrial morphology by suppressing stress-induced DRP1 activation and fission. In contrast, OA chondrocytes display elevated DRP1 activity, leading to fragmentation, ROS accumulation, and apoptosis ^49^. Pharmacological inhibition of DRP1 has been shown to mitigate mitochondrial dysfunction and cartilage damage in OA models. Interestingly, our results diverge from cardiomyocyte studies where CaMKII enhances DRP1-driven fission to support higher energy turnover ^31,50^, again suggesting tissue-specific adaptations. A limitation of our analysis is the use of MitoTracker Red CMXROS, which labels mitochondria independently of membrane potential. Although this provides a general view of morphology, it cannot distinguish between functional and depolarized mitochondria. Future studies should employ potential-sensitive dyes such as TMRE or JC-1 for more accurate assessment.

Finally, we observed that CaMKII overexpression enhances mitophagy, as indicated by increased phospho-Parkin levels. This interpretation aligns with reports of CaMKII-mediated regulation of autophagy in other contexts ^2,51^. However, mitophagy in OA appears to play dual roles: protective in maintaining mitochondrial integrity and removing severely-damaged mitochondria, yet potentially harmful when dysregulated ^52,53^. It may appear paradoxical that CaMKII simultaneously suppresses DRP1-mediated fission while promoting mitophagy, given that mitochondrial fragmentation is often a prerequisite for organelle turnover. However, our data were obtained under acute oxidative stress conditions, during which excessive DRP1 activation leads to pathological mitochondrial fragmentation, loss of membrane potential, and bioenergetic collapse. In this context, CaMKII-dependent inhibition of DRP1 likely prevents uncontrolled fission rather than abolishing the basal, localized fission events required for mitophagy. Moreover, selective mitophagy can occur in the absence of widespread fragmentation, as cells retain the ability to isolate and remove damaged mitochondrial subdomains. Thus, CaMKII may shift the balance from stress-induced catastrophic fragmentation toward a more regulated quality-control programme in which damaged mitochondria are removed through mitophagy while overall network integrity is preserved. Long-term studies will be required to determine whether CaMKII also influences mitochondrial biogenesis or remodelling under chronic stress.

Collectively, our results position CaMKII as an integrative regulator of redox and mitochondrial homeostasis in chondrocytes, coordinating NRF2 activation, respiratory support, inhibition of excessive fission, and promotion of mitophagy. While technical and contextual limitations remain, these data suggest that CaMKII contributes to mitochondrial resilience under oxidative stress and could represent a novel therapeutic target for OA as summarized in the schematic representation (Fig 6).

**Fig 6:**
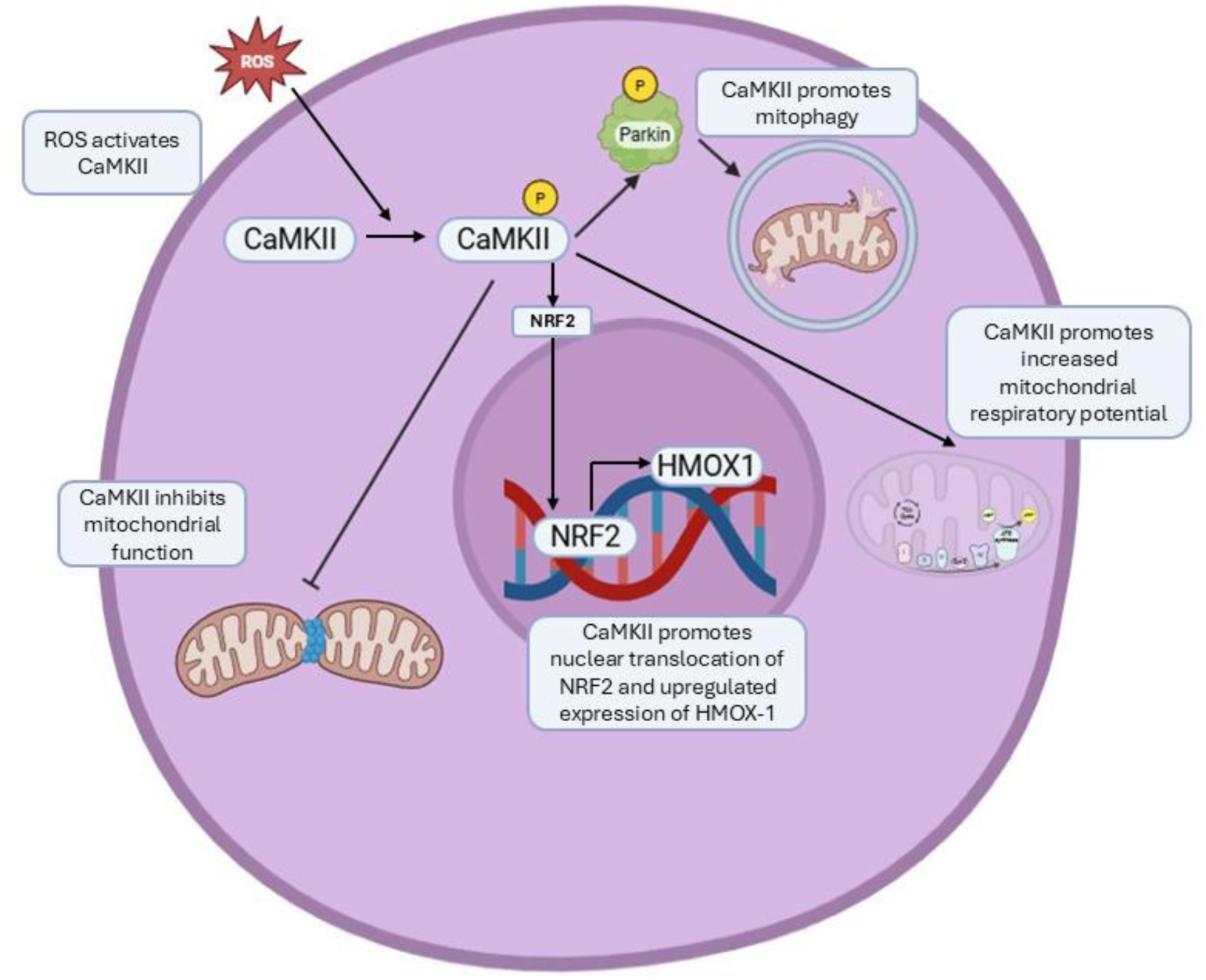
Schematic representation of the regulation of different pathways involved in cellular homeostasis by CaMKII in AHACs. ROS results in activation of CaMKII by phosphorylation in AHACs. Activated CaMKII promotes nuclear translocation of NRF2 and upregulates the expression of HMOX-1. Further activated CaMKII inhibits mitochondrial fission by downregulating the phosphorylation of DRP1 and further promotes mitophagy by upregulating the phosphorylation of Parkin. Activated CaMKII also upregulates the mitochondrial respiratory potential by increasing the spare respiratory capacity, maximum respiration, and ATP coupled respiration.

